# A global comparison of structural properties across ecological network types: the role of connectance, degree distribution and sampling inconsistencies

**DOI:** 10.1101/2024.12.04.626839

**Authors:** David García-Callejas, Elisa Thébault, Ismaël Lajaaiti, Lucas P. Martins, Louise Laux, Sonia Kéfi

## Abstract

Understanding how the structure of ecological communities varies across biotic and abiotic dimensions is a fundamental goal in ecology. This challenge is now approachable due to the increasing availability of data on community structure across the globe. Ecological communities are often defined with respect to the guilds considered and the interactions they engage in, but it is unclear whether interactions of different types respond similarly to large-scale environmental gradients. Therefore, we lack a deeper understanding of how the emergent structure of interaction networks varies across biogeographical gradients, and how this effect may change depending on their constituent interaction types. Here, using a unique dataset of 952 networks across the globe, we provide a first comparison of network structural metrics and their large-scale variability for five overarching interaction types (feeding, frugivory, herbivory, parasitism and pollination). We show that degree distribution, but not connectance alone, helps us understand the observed network structures, and this pattern is maintained across interaction types (with the partial exception of food webs). Moreover, degree distribution descriptors are generally explained by differences across studies, which represent a proxy for variability in sampling and network construction methods. Environmental factors show weaker but robust effects on network degree distribution, and food webs are generally more sensitive to changes in environmental factors than networks of other interaction types. By analysing common descriptors of the degree distributions of ecological networks, this study underscores for the first time generalities and differences across networks of different interaction types and their response to environmental and anthropogenic factors.

## Introduction

Ecological communities are emergent complex systems that vary over time, space and in response to external (e.g. environmental) factors (Levin, 1998). Understanding how the structure and dynamics of ecological communities vary with the environment is crucial to developing scientifically sound global change adaptation and mitigation programs (Harvey *et al*., 2017). However, this is a challenging task due to the myriad of interactions and potential confounding factors that may influence these relationships (Tylianakis & Morris, 2017). Conceptualising ecological communities as networks composed of species and their interactions provides a powerful tool to study community structure and its drivers, as it allows us to separate these components and their potential feedbacks (Vázquez *et al*., 2009).

The variability in network structure across space can be decomposed into the variability at-tributed to species turnover on the one hand, and the variability attributed to interaction turnover on the other, i.e. the change in presence and strength of biotic interactions between species (Poisot *et al*., 2012). Although much is known about how species are distributed in space, our knowledge of how network structure changes across environmental gradients is still unfolding (Chamberlain *et al*., 2014; Early & Keith, 2019). Recent studies have shown that biotic interactions are expected to vary with environmental factors (Poisot *et al*., 2017) or geographical proxies such as latitude or altitude (Roslin *et al*., 2017; Zvereva & Kozlov, 2021, 2022), but the ecological processes behind these patterns and the emerging consequences for community structure and dynamics have not yet been sufficiently explored.

Different types of interactions can show specific responses in their frequency and intensity to changes in biotic or abiotic drivers. Latitudinal effects on interaction strength, for example, are different for herbivory and carnivory when compared to parasitism, and further depend on the metabolism of the involved species (e.g. ectotherms or endotherms) (Zvereva & Kozlov, 2021). Moreover, species-specific responses to temperature variability may trigger phenological mismatches between interacting species, which are more prevalent in species with strong responses to temperature cues, such as flowering plants and their insect pollinators (Gérard *et al*., 2020). Likewise, temperature is directly related to metabolic rates, which influences e.g. movement speeds in ectotherms, with direct consequences for the frequency of interactions between individuals (Early & Keith, 2019).

How these pairwise effects translate into the structure and dynamics of entire communities, and whether there exist differences across interaction types is not yet fully resolved. Thébault & Fontaine (2010) showed that mutualistic bipartite networks tend to be more nested and less modular than trophic networks, and these patterns have a positive effect on the stability of these systems. Further studies have challenged these conclusions, by showing that bipartite binary mutualistic and antagonistic newtorks are impossible to tell apart only based on their structure (Michalska-Smith & Allesina, 2019). Including environmental context (Song & Saavedra, 2020) or meso-scale structural information (Pichon *et al*., 2024) improves the structural separability of mutualistic and antagonistic networks, as well as explicitly accounting for interaction type (Pichon *et al*., 2024). Together, these advances suggest that network structure across different types of interactions is difficult to distinguish, although meso-scale patterns and external factors may help us discern this variability.

Network structure is not a static property of ecological communities, however. Different descriptors of network structure vary in response to external factors (Tylianakis & Morris, 2017), including environmental, anthropogenic, or other biotic factors (e.g. species richness). Besides the unresolved question of whether there exist fundamental differences between the structure of different types of interaction networks, a related issue is whether different types of networks respond in similar or different ways to variations in such external factors. Evidence regarding this question is elusive, as studies often suggest contrasting relationships between network properties and environmental factors. For example, several studies have analysed the relationship of nestedness, a network structural property (Mariani *et al*., 2019), with climatic seasonality at biogeographical scales, finding negative (Takemoto & Kajihara, 2016), positive (Song *et al*., 2017), or no relation-ship (Brimacombe *et al*., 2022) for plant-pollination networks, and no relationship for frugivory or host-parasite networks (Brimacombe *et al*., 2022). Overall, relationships between large-scale environmental variation and network structure have been studied generally independently across different network types, with contrasting aims and methodologies, precluding generalisations.

Combining these lines of evidence, there is an emerging consensus that local community structures potentially vary across interaction types. In parallel, pairwise interactions of different types display different trends in their spatial variation and response to external factors. Therefore, we might expect the variation in community structure across large spatial gradients, and in response to external factors, to vary across different interaction types. Alternatively, some factors structuring ecological networks across space may be similar across interaction types. For example, increasing richness towards the tropics may influence fundamental structural network patterns, such as connectance, regardless of interaction type (Dallas & Jordano, 2021; Gibert & Wieczynski, 2021). Looming over these considerations is the challenge of finding ecological signals over data on interaction networks from a large spatial extent, with vastly different collection methodologies and initially gathered for potentially different research questions (Brimacombe *et al*., 2023). Under that light, assessing how these inconsistencies affect ecological networks of different interaction types is also paramount. Here, we undertake a comparative analysis of structural network metrics at a global scale by using an unprecedented compilation of 952 ecological networks including five overarching interaction types (pollination, frugivory, herbivory, parasitism, and food webs). We first evaluate whether fundamental network properties (connectance and degree distribution) explain the observed structural patterns in the networks compiled, and then evaluate the biogeographical factors and sampling biases influencing the degree distribution of ecological networks across interaction types.

## Methods

### Network datasets

We compiled ecological network data across the globe, considering undirected binary interactions, uni- or bipartite networks, and with associated spatial coordinates that we could link to a geo-graphical location. We combined networks from open repositories (mangal (Poisot *et al*., 2016), Web of Life, GATEWAy (Brose *et al*., 2019)) and from published studies (Boscolo *et al*., 2023; Dalsgaard *et al*., 2021; Fricke & Svenning, 2020; Martins *et al*., 2022; Parra *et al*., 2022) as well as additional datasets collected by the authors. We classified each network according to its inter-action type (pollination, frugivory, herbivory, parasitism, or predation) and topology (unipartite or bipartite). The first set comprised 4341 ecological networks, subsequently filtered according to the following criteria: First, we selected networks with > 20 species for unipartite networks, or alternatively, > 10 species in each of the groups in bipartite networks. Second, we selected networks that were not fully connected, i.e. where not all species interacted with one another, as these usually represent small, finer-scale studies where community structure is not the focus. With these constraints, we aimed to select networks that represented at least moderately diverse communities, with sufficient richness so that structural metrics could be reliably computed and would be ecologically meaningful. The final dataset is, to our knowledge, the most comprehensive compilation of binary networks across interaction types so far: it comprises 61 frugivory, 444 pollination, 98 herbivory and 39 parasitism networks, as well as 310 food webs, for a total of 952 networks distributed across all continents (Fig. 1). Among those, 642 are bipartite and 310 are unipartite food webs.

**Fig. 1:**
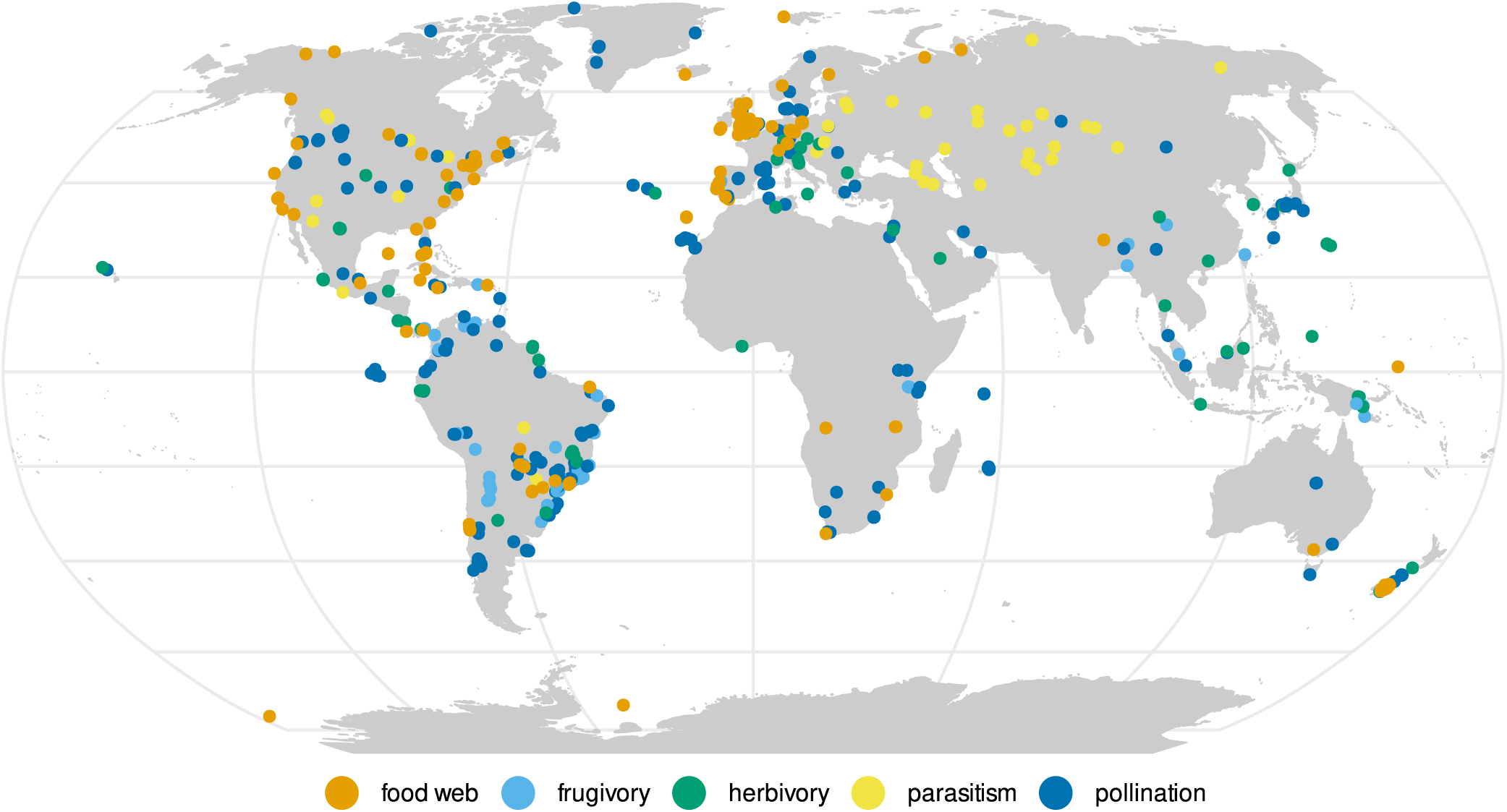
Map displaying the spatial location of the ecological networks included in this study. N = 952 networks, of which 310 food webs, 61 frugivory, 98 herbivory, 39 parasitism, and 444 pollination networks.

### Structural network metrics

We calculated a suite of structural metrics for every network in our dataset. More specifically, we obtained the interaction overlap (García-Callejas *et al*., 2023), nestedness (Delmas *et al*., 2018; Mariani *et al*., 2019), modularity using two different algorithms (infomap (Farage *et al*., 2021) and an algorithm based on betweenness centrality of nodes (Newman & Girvan, 2004)), and eigenvector centralisation, a metric representing the eigenvector centrality aggregated at the network level (Delmas *et al*., 2018). These metrics represent a variety of structural properties of networks and their interpretation is conceptually similar in uni- and bipartite networks. Furthermore, all these metrics can be applied to networks with more than one component, i.e. networks that consist of more than one disconnected subnetworks or components, where no links connect nodes in different components. This was the case for 399 networks.

The degree distribution of a network lists the number of links of each node in the network. Like any other distribution, degree distributions cannot be fully reduced to a one-dimensional metric. Hence, we obtained the first four statistical moments of the degree distribution of each network in our data: mean, variance (or standard deviation), skewness, and kurtosis. Mean and standard deviation were positively correlated across the dataset (Pearson’s *ρ* = 0.76, p *<* 0.05, Fig. S2), as were skewness and kurtosis (*ρ* = 0.902, p *<* 0.05, Fig. S2). We therefore consider in the following analyses both the mean and skewness of our network’s degree distributions, discarding their kurtosis. Further, as standard deviation is an important descriptor of statistical distributions, we also include it despite its relatively strong correlation with the mean of the degree distributions.

### Null models

To be properly interpreted, it is important to assess the statistical significance of the structural metrics obtained, i.e. whether the value of a given metric for a given network can be expected solely by looking at general network properties such as the number of nodes or links in the network, or whether there are ecological processes potentially driving it (Delmas *et al*., 2018). To do so, for each network, we generated a set of 999 randomised realisations considering three different null models with increasing constraints. The first null model constrained connectance values; the second and third null models constrained, in addition to connectance, the full degree distribution of the networks. The difference between our second and third null models is whether this constraint is strictly enforced in each null realisation. In the second null model (hereafter *soft-constrained degree distribution*), the degree distribution of each null realisation is allowed to vary slightly from the observed degree distribution, while maintaining the exact same degree distribution as the original network *on average* (Caruso *et al*., 2022). The third null model (hereafter *hard-constrained degree distribution*) strictly enforced the same degree distribution of the observed network in each null realisation. This approach allowed us to test whether a given structural property of an observed network could be explained simply by knowing the network’s connectance, its approximated degree distribution, or its exact degree distribution. We implemented the connectance null model with the shuffle.web algorithm of the bipartite R package v2.20 (Dormann *et al*., 2009), which is based on the Patefield algorithm (Blüthgen & Staab, 2024) but does not constrain marginal sums, i.e. species degrees. For the soft-constrained degree distribution null model, we used the NEMtropy Python package (Vallarano *et al*., 2021), and for the hard-constrained degree distribution null model, the curveball algorithm (Strona *et al*., 2014) implemented in the vegan R package v2.6-8 (Oksanen *et al*., 2024).

For each network, we compared the values of each metric in the observed network with the distribution of values obtained in the randomised network realisations. Then, for each metric, we obtained an associated z-score or standardised effect size (Ford & Roberts, 2019), that quantifies the difference between the observed value and the distribution of null realisations:

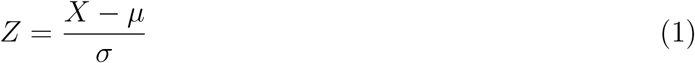

where the z-score *Z* is a function of the observed value of a metric in a network (*X*), the mean of the metric value for the null realisations (*µ*) and its standard deviation (*σ*). Values of *Z >* 2 or *Z <* −2 indicate statistically significant differences between them.

### Biogeographic determinants of degree distribution descriptors

Null models constraining degree distribution generally reproduced the structural characteristics of the observed networks, whereas the null model constraining only connectance did not (see Results). Therefore, we turned to analyse the variability of degree distribution descriptors (mean, standard deviation, and skewness) across interaction types globally, and how they were influenced by environmental factors. We obtained WorldClim environmental data at 1km resolution (Fick & Hijmans, 2017) and used annual mean temperature and precipitation, as well as their seasonality indices, as independent variables in our models. Spearman’s correlations between these variables in the spatial locations of our networks (Fig. 1) were all *<* 0.5 except between annual mean temperature and temperature seasonality (*ρ* = −0.78, see Fig. S1). We nevertheless decided to keep temperature seasonality alongside the other environmental variables in the following analyses because it has been extensively explored in previous studies of biogeographical network structure, and excluding it did not improve goodness-of-fit of the statistical models detailed below.

We fitted linear mixed-effects models for each interaction type, taking each of the three degree distribution descriptors as response variable: a) mean degree distribution, b) standard deviation, and c) skewness. For these 5 ∗ 3 = 15 models, we only included networks with available spatial information for all variables. We considered the following predictors: annual mean temperature, temperature seasonality, annual mean precipitation, and precipitation seasonality, as defined above. All independent variables were scaled before performing the analysis. In addition, variability in network structure is known to be highly driven by the variability across individual studies, both in bipartite networks (Brimacombe *et al*., 2023) and unipartite food webs (Brimacombe *et al*., 2024). We therefore included study as a random intercept effect in our models. Because many studies in our dataset - and therefore levels of the random effect-consist of a single network (Brimacombe *et al*., 2024), we filtered the dataset to account only for studies with 3 or more networks. Thus, our final datasets for the statistical analyses consisted of 258 food webs from 10 studies, 13 frugivory networks from 2 studies, 33 herbivory networks from 4 studies, 30 parasitism networks from 2 studies, and 276 pollination networks from 36 studies, totalling 610 networks. Including studies with one or two networks in the datasets did not modify the qualitative trends presented here. We implemented all models in R v4.3.2 (R Core Team, 2023) using the package glmmTMB v1.1.9 (Brooks *et al*., 2017), and checked model fits with the DHARMa package v0.4.6 (Hartig, 2022). We obtained the associated *R*^2^ values associated to the fixed effects of environmental variables (marginal *R*^2^) and to the fixed and random effects together (conditional *R*^2^) following Nakagawa *et al*. (2017) as implemented in the performance package v0.11 (Lüdecke *et al*., 2021). The difference between the conditional and marginal *R*^2^ is indicative therefore of the variance explained by the random effect (individual study) compared to that explained by the fixed effects only (environmental variables).

In addition to these main statistical models, we generated ancillary linear mixed-effects models to test for differences in 1) network raw structural metrics across interaction types, 2) z-score distributions across interaction types, and 3) degree distribution descriptors across interaction types. In these models, we tested for pairwise differences between interaction types with post-hoc Tukey tests, using the emmeans package v1.10.5 (Lenth, 2024).

## Results

### Structural metrics across interaction types

The raw values of the network metrics analysed tend to differ across interaction types (top row of Fig. 2). In particular, for four out of the five metrics, networks can be categorised in four groups with significantly different metric values: food webs (group a) in Fig. 2; frugivory and parasitism networks (group b); herbivory (group c); and pollination (group d). Food webs showed the lowest modularity and eigenvector centralisation, and intermediate nestedness. Herbivory networks, fol-lowed by pollination networks, were the most modular and centralised, and the least nested; other networks showed intermediate values. Connectance was positively correlated with interaction over-lap, nestedness, and negatively correlated with modularity and centralisation metrics (middle row of Fig. 2, see Fig. S3 for Spearman correlation values). The correlations between metrics and the mean of the degree distribution (bottom row of Fig. 2) were of similar sign but generally weaker, especially for food webs.

**Fig. 2:**
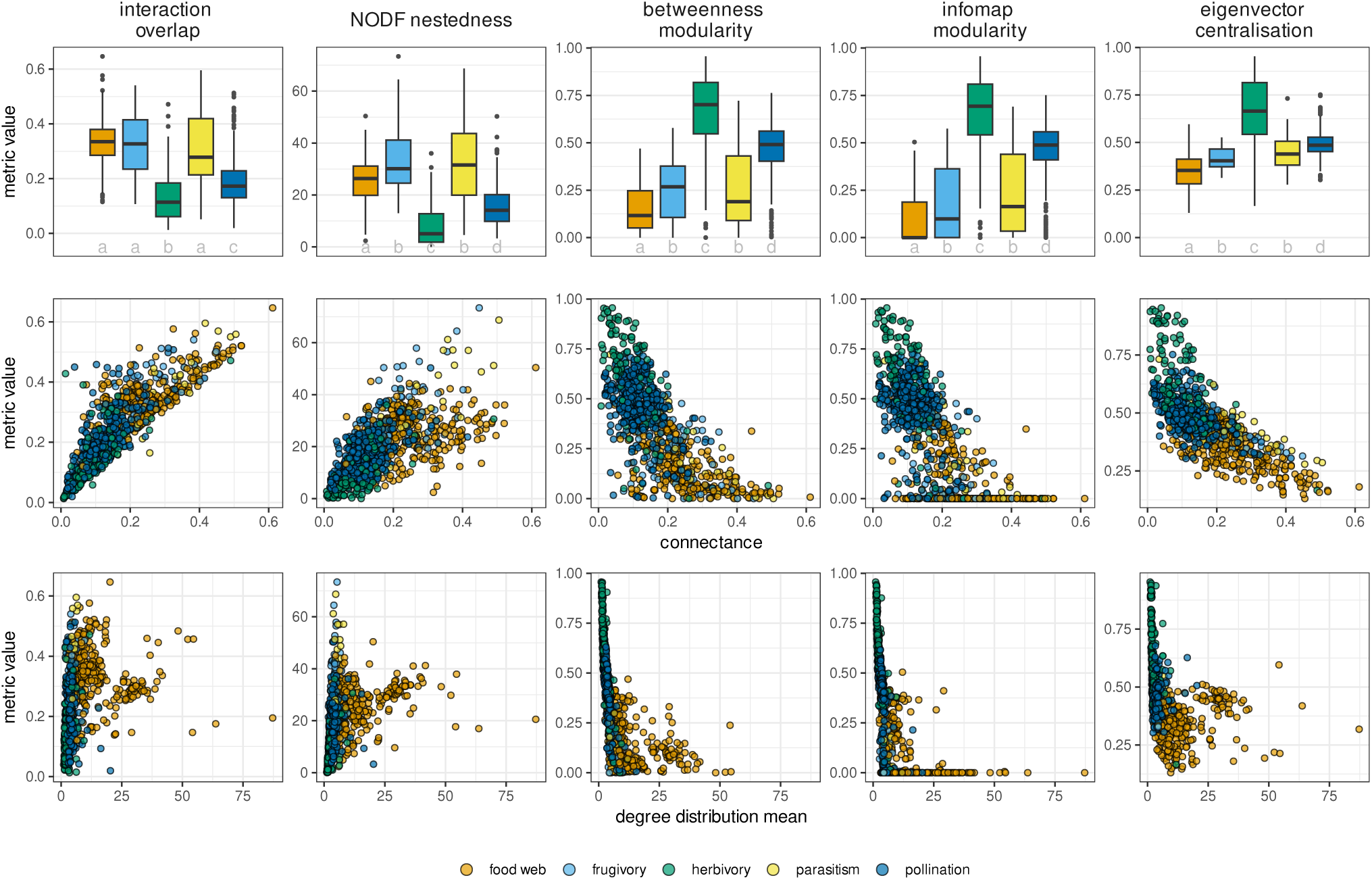
Structural metric values for ecological networks (top row), and their relationship with network connectance (middle row) and degree distribution mean (bottom row). For all plots, N(food webs) = 310, N(frugivory) = 61, N(herbivory) = 98, N(parasitism) = 39, N(pollination) = 432. Letters at the bottom of boxplots correspond to statistically significant differences between distributions obtained from a generalised linear mixed model with interaction type as fixed effect and study as random intercept. In the boxplots, the horizontal black line represents the median, the left and right hinges correspond to the 25th and 75th percentiles, and the vertical lines extend to the largest/smallest value up to 1.5 times the interquartile range (distance between 25th and 75th percentiles). Single points are observations outside of 1.5 times the interquantile range.

### Null model analysis

Despite the correlations observed between network metrics, connectance, and degree distribution mean (Fig. 2, Fig. S3), null models constraining connectance were not able to recover the observed metric values. However, null models constraining the exact (and sometimes approximate) degree distribution showed values similar to the observed networks (Fig. 3). For the null model constrain-ing connectance, our results show 1) highly variable z-scores for all network types, and 2) observed values much larger than those generated by the null models for interaction overlap, nestedness, and eigenvector centralisation, with modularity values closer to the observed ones. The second null model, softly constraining degree distribution, better approximated the observed metric values, but the observed betweenness and infomap modularity tended to be higher than in the null realisations. Exactly constraining degree distribution, in turn, generated null network realisations that mostly showed similar values to the observed networks, with the notable exception of food webs. As shown in the bottom row of Fig. 3, food web metrics except nestedness were generally higher in the observed than null networks, due to their z-scores being, on average, near or above statistical significance (z-score > 2). This finding reveals that connectance by itself is not enough to explain observed network structure; in most cases, the approximate or exact degree distribution is needed, but for food webs, even this information does not fully capture their structural patterns.

**Fig. 3:**
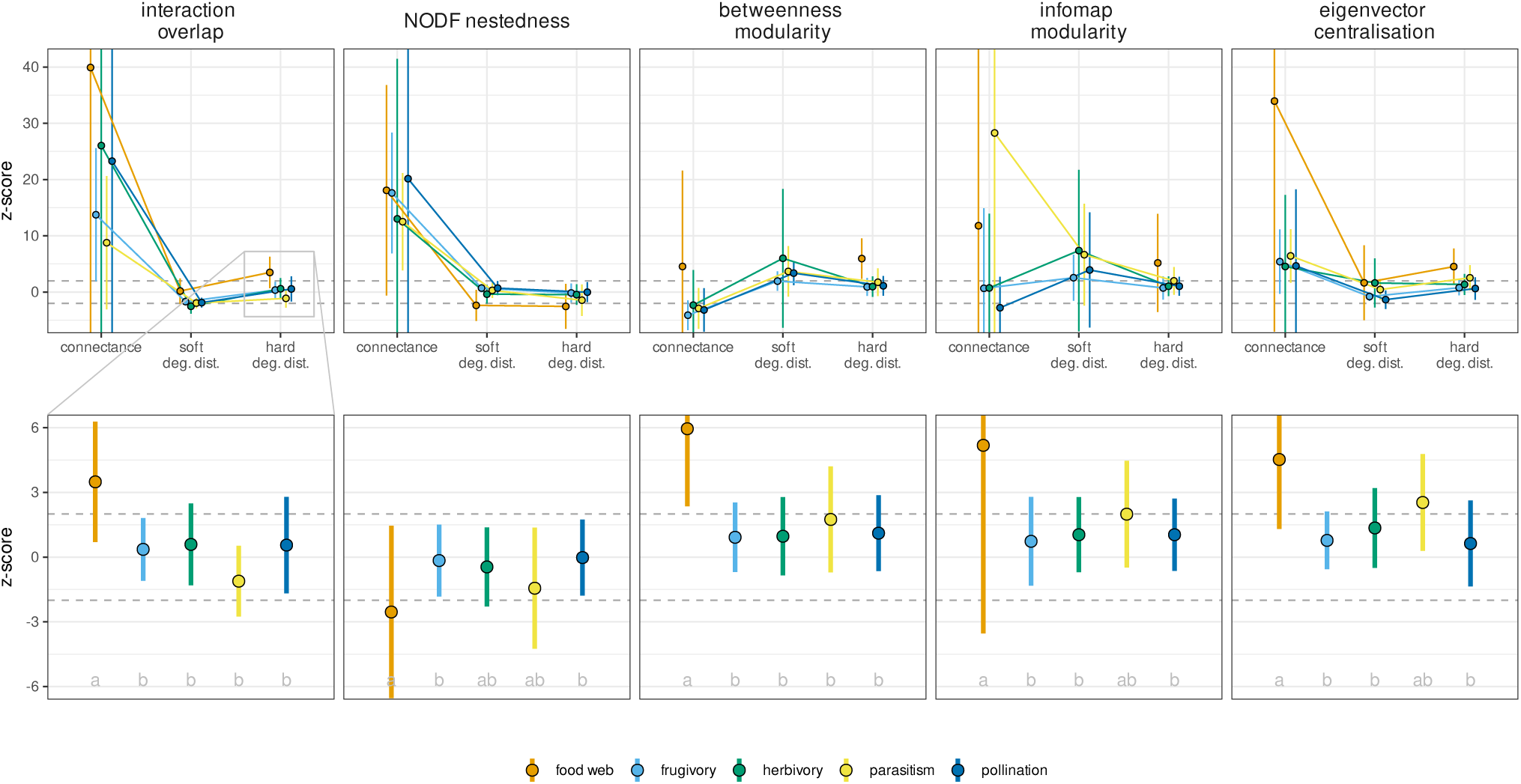
Z-scores of three null models (x-axis of top panels) for different structural metrics (columns) and interaction types (colors). Dashed horizontal lines in both panels represent the limits of statistical significance for z-scores, whereby values outside them represent statistically significant differences between observed and null values. In the top-row panels, the z-scores of the three null models are shown sequentially, with the x axis indicating null models from least to most constrained. Points represent mean values, and lines extend to one standard deviation from the mean. In these panels, vertical axis is limited to 40 for visibility. Two points not shown in the figure are above this threshold: betweenness modularity of food webs for the soft-constrained degree distribution null model, with a value of 56.2; and infomap modularity of food webs for the soft-constrained degree distribution null model, with an extreme value of 2.57 · 10^13^. The lower panels show in more detail the z-scores of the hard-constrained degree distribution null model. Letters at the bottom of these panels correspond to statistically significant differences between interaction types obtained from a generalised linear mixed model with interaction type as fixed effect and study as random intercept.

### Biogeographical determinants of degree distributions

The findings of Fig. 3 highlight the importance of degree distributions in driving network properties, and hence raise the questions of how degree distributions vary across networks of different interaction types and what factors are shaping these patterns across large spatial scales. First, degree distribution descriptors in our dataset tended to differentiate food webs from other interaction types, with food webs showing higher mean and standard deviation of the degree distribution, and lower skewness and kurtosis (Fig. 4). Interestingly, degree distributions of bipartite networks (frugivory, herbivory, parasitism, and pollination) were similar on average, without consistent statistical differences across interaction types.

**Fig. 4:**
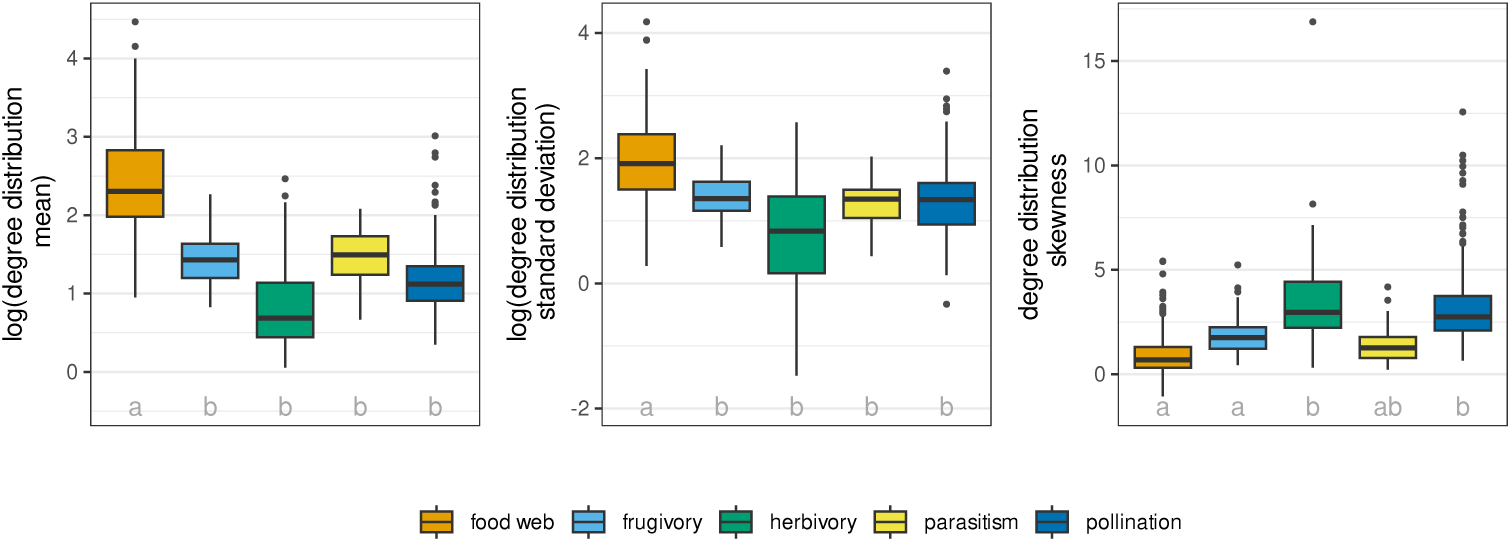
Statistical moments (mean, standard deviation, and skewness) of the degree distribution of the 952 networks analysed, differentiated by interaction type. Food webs are all unipartite networks; all other interaction types are bipartite. In the vertical axis, values are log-transformed for visibility, except for skewness (right panel), that can take negative values. Letters at the bottom of each panel represent statistically significant groupings obtained from a generalised linear mixed model with interaction type as fixed effect and study as random intercept. Boxplot features as in Fig. 2.

The variability in these metrics was in general not statistically related to environmental variables (Fig. 5). In particular, only food webs and herbivory networks showed statistically significant relationships between degree distribution metrics and environmental variables: the mean and standard deviation of degree distributions in food webs were negativaly related both to temperature seasonality and annual mean precipitation; skewness was negatively related to mean temperature and its seasonality, and positively related to annual mean precipitation. In herbivory networks, the standard deviation of their degree distributions was related positively to annual mean temperature and negatively to annual mean precipitation, whereas their skewness was negatively related to temperature seasonality. No other interaction type showed statistically significant relationships with these large-scale environmental factors.

**Fig. 5:**
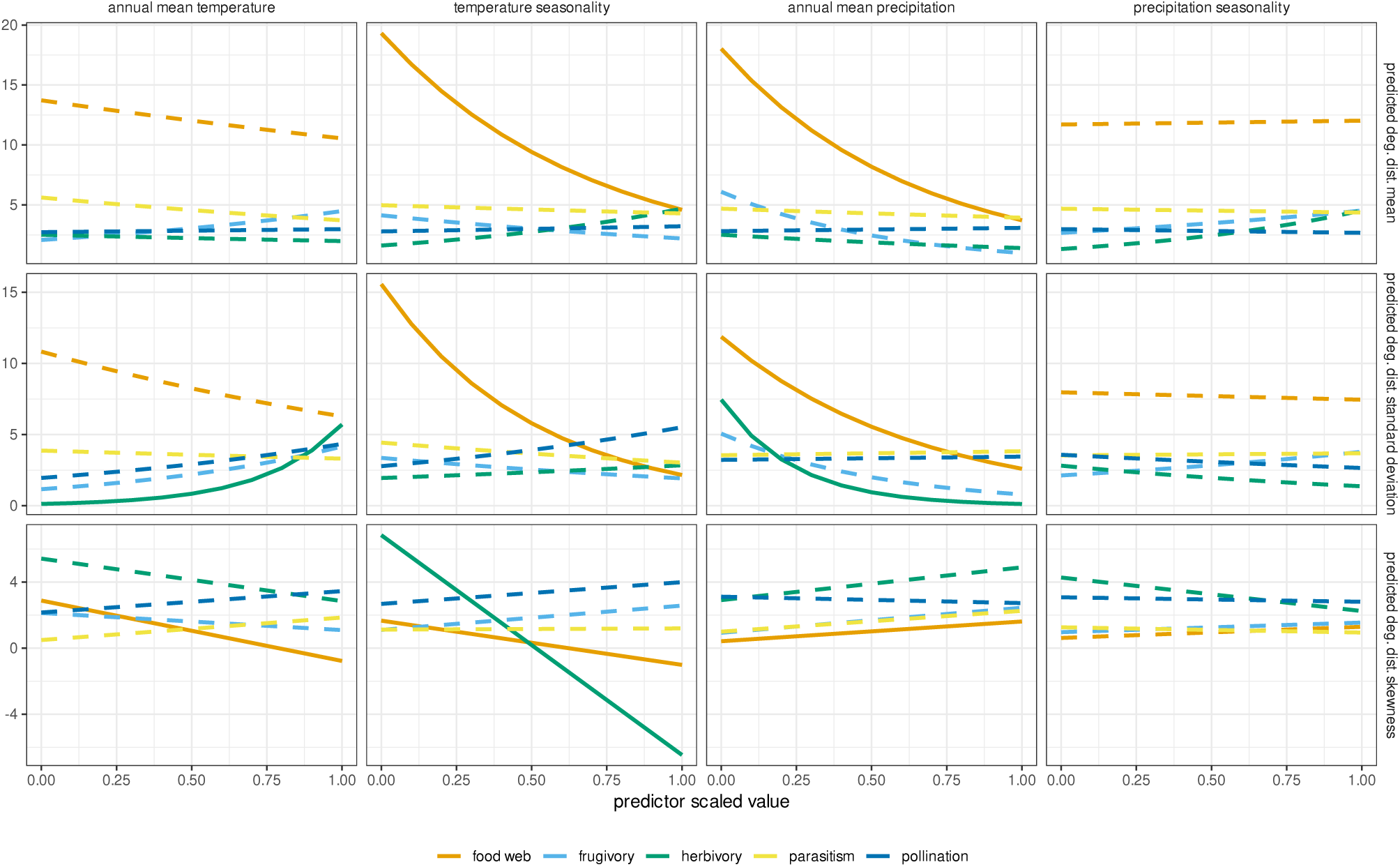
Marginal effects of each scaled predictor variable on degree distribution mean (top row), standard deviation (middle row), and skewness (bottom row), for each interaction type. Effects of each predictor are obtained by setting non-focal predictors to their mean, using the package ggeffects v1.6 (Lüdecke, 2018). Solid lines represent statistically significant effects, dashed lines non-significant ones. Confidence intervals are not shown for visibility, as they are highly overlapping in most cases. Effects for degree distribution mean and variance may show non-linear patterns because the response variable is log-transformed in the original model, and predicted values are back-transformed to the original response scale.

Models with or without including study as a random effect showed generally similar qualitative trends in terms of effects direction and magnitude (Fig. S4), so hereafter we discuss models including it as they showed consistently lower AIC scores. Accordingly, when included, the study random effect accounted for a much higher percentage of explained variance than the fixed effects (Table 1, Fig. 6) across all interaction types with sufficient data (food webs, pollination networks, and partly herbivory networks). This suggests that variability across individual studies is by far the largest source of variability in the degree distributions of our dataset, but the fact that the trends remain similar with or without considering across-study variability suggests that the environmental effects modelled are robust to it.

**Fig. 6:**
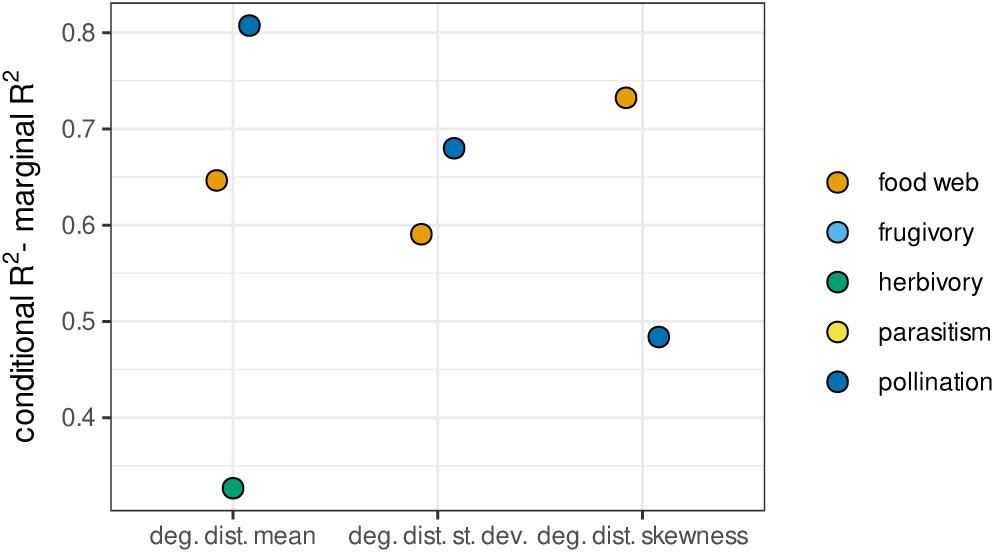
Difference between conditional and marginal R^2^ for each model, from the raw data of Table 1. Only differences for models where we obtained both conditional and marginal R^2^ values are shown.

**Table 1:** *R*^2^ values of the linear mixed models relating degree distribution descriptors (log-transformed mean, log-transformed standard deviation, and skewness) with environmental variables and study as random factor. Marginal *R*^2^ values capture the variance explained by fixed effects, and conditional *R*^2^ the variance explained by fixed and random effects together. Conditional *R*^2^ values for some models were not obtained due to the small number of random factor levels (i.e. small number of different studies).

| Interaction type | $R^2$ type | response | $R^2$ value |
| --- | --- | --- | --- |
| food web | conditional | mean | 0.81 |
| food web | conditional | st. dev. | 0.83 |
| food web | conditional | skewness | 0.78 |
| food web | marginal | mean | 0.16 |
| food web | marginal | st. dev. | 0.24 |
| food web | marginal | skewness | 0.05 |
| frugivory | conditional | mean | - |
| frugivory | conditional | st. dev. | - |
| frugivory | conditional | skewness | - |
| frugivory | marginal | mean | 0.57 |
| frugivory | marginal | st. dev. | 0.65 |
| frugivory | marginal | skewness | 0.03 |
| herbivory | conditional | mean | 0.96 |
| herbivory | conditional | st. dev. | - |
| herbivory | conditional | skewness | - |
| herbivory | marginal | mean | 0.63 |
| herbivory | marginal | st. dev. | 0.85 |
| herbivory | marginal | skewness | 0.60 |
| parasitism | conditional | mean | - |
| parasitism | conditional | st. dev. | - |
| parasitism | conditional | skewness | - |
| parasitism | marginal | mean | 0.04 |
| parasitism | marginal | st. dev. | 0.04 |
| parasitism | marginal | skewness | 0.20 |
| pollination | conditional | mean | 0.81 |
| pollination | conditional | st. dev. | 0.72 |
| pollination | conditional | skewness | 0.50 |
| pollination | marginal | mean | 0.01 |
| pollination | marginal | st. dev. | 0.04 |
| pollination | marginal | skewness | 0.02 |

## Discussion

Biotic interactions vary across biogeographical scales in response to factors such as temperature (Dell *et al*., 2013), precipitation and water availability (Walter, 2018), or human pressure (Sebastián-González *et al*., 2015). While recent studies have started to explore how environmental, sampling, or anthropogenic factors influence the structure of ecological networks across large spatial scales (Brimacombe *et al*., 2022; Martins *et al*., 2022), it is still unclear whether different types of biotic interactions show similar or different patterns in their variation. Here, we have shown that the structure of ecological networks across large spatial scales displays both commonalities and differences across interaction types, that their degree distribution is a fundamental property to disentangle this variability, and that sampling inconsistencies across studies generally overcome environmental influences in shaping network structure across large spatial scales.

The degree distribution of an ecological network informs about the number of interactions that each species has with the rest of the species pool. As such, together with connectance (Poisot & Gravel, 2014), several secondary properties can potentially emerge from it. In our comparative analyses, we showed that structural properties of ecological networks cannot be explained simply by their connectance; rather, knowing the degree distribution is necessary to reproduce the observed properties. In most cases, an approximate degree distribution in which the null realisations are allowed to deviate slightly from the observed degree distribution accurately recovered structural properties. Such soft-constrained null models may be conceptually more fit to the analysis of ecological networks than hard-constrained ones, as it explicitly integrates uncertainty in the observed network structure (Caruso *et al*., 2022; Neal *et al*., 2024). Robustly quantifying degree distribution may be particularly challenging for communities with a high prevalence of rare and specialist species. Rarity is, however, common in ecological communities (Enquist *et al*., 2019), and empirical studies have repeatedly shown that, in order to estimate interactions accurately, it is often necessary to combine several sampling methods (Bosch *et al*., 2009; Quintero *et al*., 2022) to avoid potential biases by which abundance alone would drive the number of interactions detected (Blüthgen & Staab, 2024). Under this light, degree distributions such as the ones analysed here are subject to substantial sources of uncertainty: soft-constrained null models may be more conceptually aligned with this type of empirical data.

These considerations apply both to unipartite and bipartite networks of different interaction types, and to binary and weighted ones. However, unipartite food webs tended to show larger differences than bipartite networks between the observed networks and our two null models fixing degree distributions (Fig. 3). To understand this, we may combine a structural and an ecological perspective. First, from a purely structural point of view, the number of potential interaction partners for a given species is larger in a unipartite network of richness *S* than in a bipartite network, simply because in bipartite networks no within-group links are allowed. This leaves a larger set of potential networks that comply with a given degree distribution. Second, from an ecological point of view, a variable fraction of these randomised interactions would effectively be forbidden interactions (Jordano, 2016). Food webs in particular are highly structured by trait matching, including body size relationships (Brose *et al*., 2019), which implies that a potentially relevant fraction of the links assigned in food web null models may correspond to forbidden interactions, generating network structures that deviate from the observed ones. Therefore, null models that explicitly incorporate trait-matching constraints in food webs, in addition to constraining degree distributions (see e.g. McLeod & Leroux (2021)), are likely to better recover the structural patterns of empirical food webs. For bipartite networks of different types, we observed a comparatively lower relevance of trait matching on top of degree distribution constraints - *on average*. This does not preclude the importance of trait matching in specific bipartite communities, as empirically observed e.g. in some frugivory networks (Sebastián-González *et al*., 2017; Vizentin-Bugoni *et al*., 2014), but rather may be useful as a baseline expectation from which ecologically relevant differences may be drawn.

To analyse degree distributions at large scales, we looked at their mean, standard deviation, and skewness. Previous studies already showed that variability across individual studies can obscure or override structure-environment relationships in ecological network analysis (Brimacombe *et al*., 2024, 2023). Here we quantified for the first time this influence on degree distribution descriptors, and found that the study-level effect accounted for much of the variance captured by our statistical models (Table 1, Fig. 6).

Despite this importance, we found a few consistent relationships between environmental factors and degree distribution descriptors, again with food webs showing different trends than other network types. Temperature seasonality and mean annual precipitation were negatively related to the number of links per species in food webs and their standard deviation, but the effect on degree distribution skewness was opposite: negative for temperature seasonality and positive for mean annual precipitation. Unexpectedly, besides food webs and herbivory networks, no other interaction type showed significant relationships between degree distribution descriptors and environmental factors. Together, these results suggest a higher environmental sensitivity of food web structure to large-scale environmental gradients than that of other interaction types. Previous studies have empirically demonstrated higher predation intensity with increasing temperature (Roslin *et al*., 2017; Zvereva & Kozlov, 2021), via an increase in predator attack rates (Tylianakis & Morris, 2017). Whether these direct effects of temperature in pairwise interactions translate into food web structural metrics is contested: Gibert (2019) found that connectance increased with increasing temperature, through a concomitant reduction in the number of basal species, whereas Gauzens *et al*. (2020) found no strong environment-structure relationships in intertidal food webs across the globe. These inconsistent results between previous studies and our findings may be due to several reasons, including the habitat type(s) of the food webs analysed. In particular, we pooled together food webs from marine (153), freshwater (91), and terrestrial (66) ecosystems, whose species and interactions may be subject to different eco-evolutionary drivers and responses to environmental factors. The absence of consistent degree distribution-environment relationship for other network types may arise from intrinsic ecological patterns, such that direct effects on species composition or metabolism may not be evident at the network level, and/or may be due to limitations of the data and analyses, e.g. in the number of networks included, binary as opposed to quantitative networks, or spatial coverage. We provide open access to our harmonisation procedure and the full dataset (see Data and code availability), hoping to spur further analyses and additions to our insights.

Overall, we have shown that currently available global data on ecological network structure suggests that 1) ecological networks of different interaction types are well characterised, in general, by their degree distribution, with the possible exception of food webs (which potentially require further trait-based constraints to understand their observed structure); 2) degree distribution descriptors of all interaction types are more strongly influenced by variations across individual studies than by large-scale environmental factors, reinforcing the importance of local contexts for determining the structure of ecological networks; and that 3) food webs, and to a lesser extent herbivory networks, are in general more sensitive to varations in environmental factors than other types of interaction networks. These insights highlight that focusing on fundamental network properties such as degree distribution may bring novel perspectives in the biogeographical study of ecological networks, but the challenge remains on obtaining ecologically relevant signals from highly heterogeneous data.

