## Supplementary Material for "A global comparison of structural properties across ecological network types: the role of connectance, degree distribution and sampling inconsistencies"

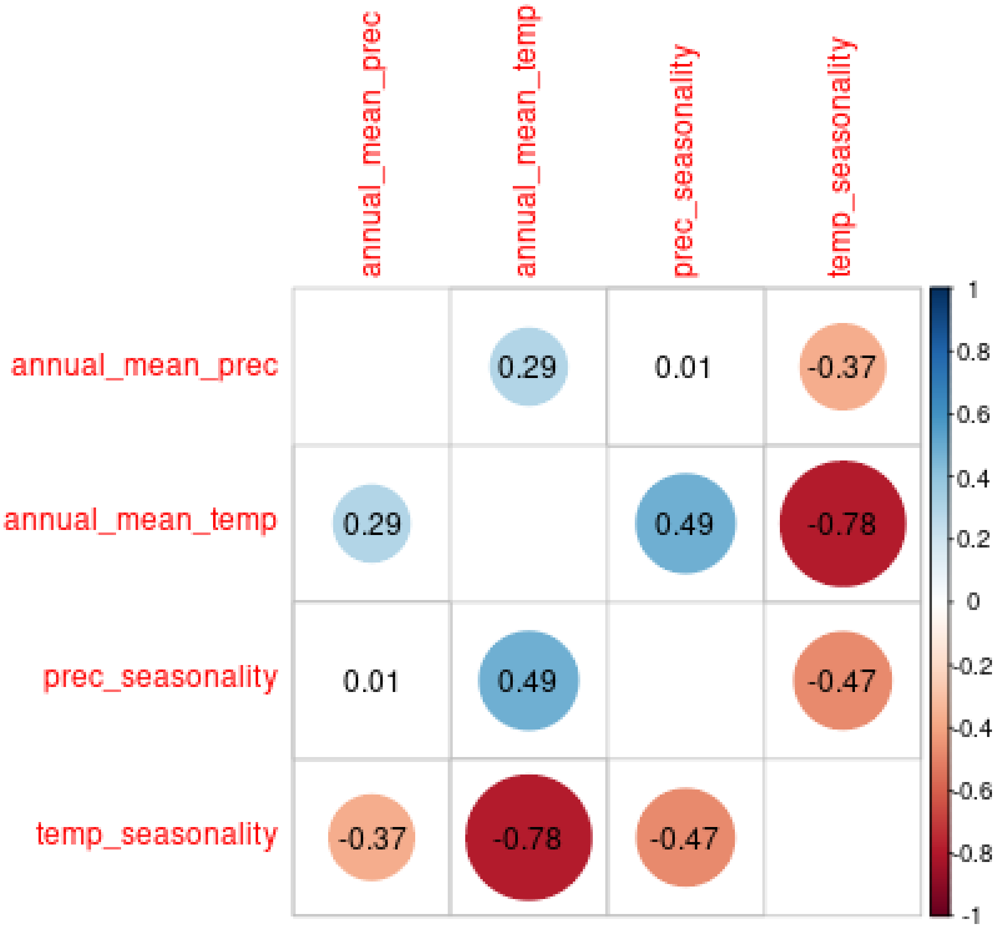


Fig. S1: Spearman correlations between environmental variables obtained for each network location (see Fig. 1 of main text). Statistically significant correlations are displayed with a colored circle.

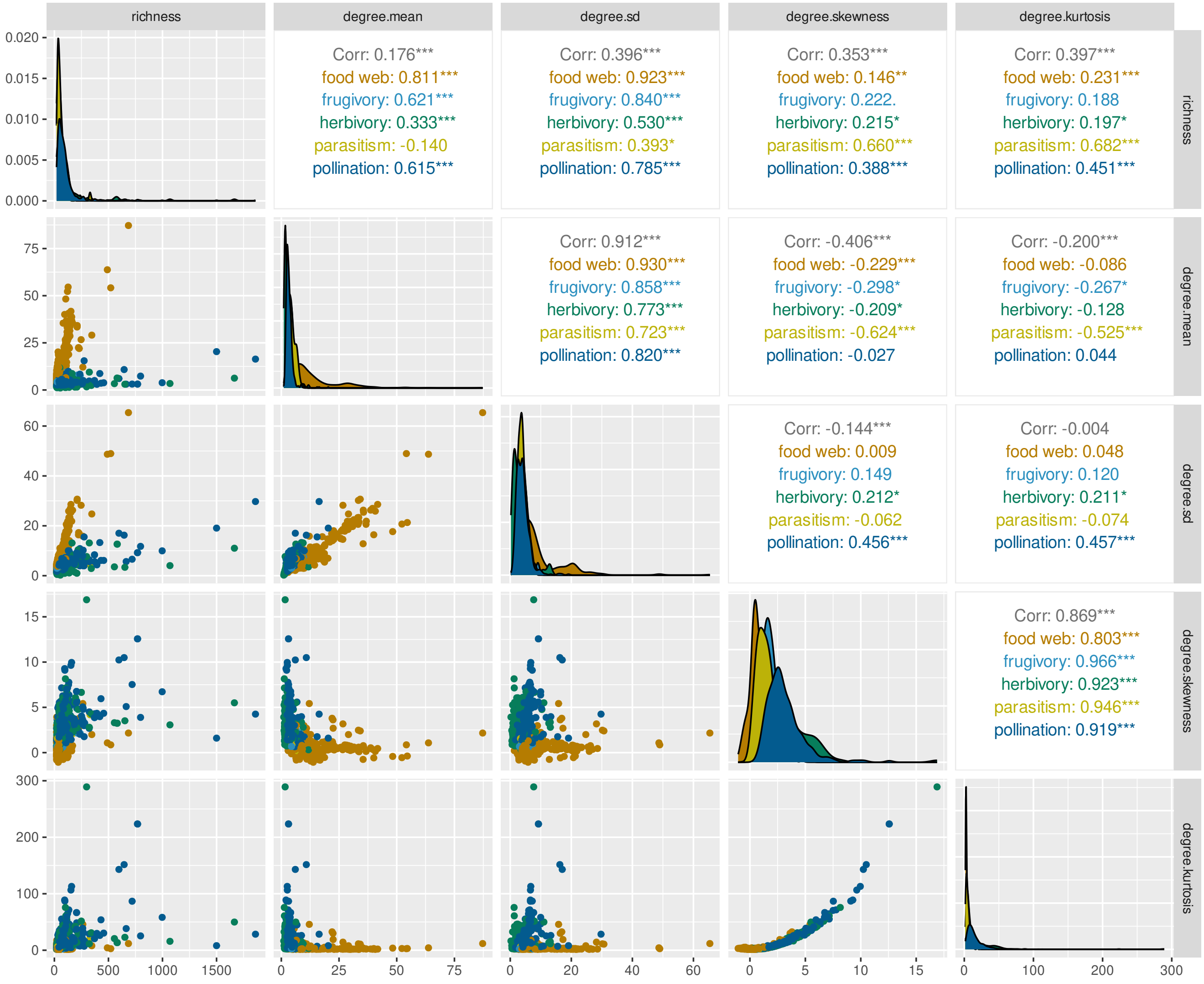


Fig. S2: Density plots and correlations between the degree distribution descriptors.

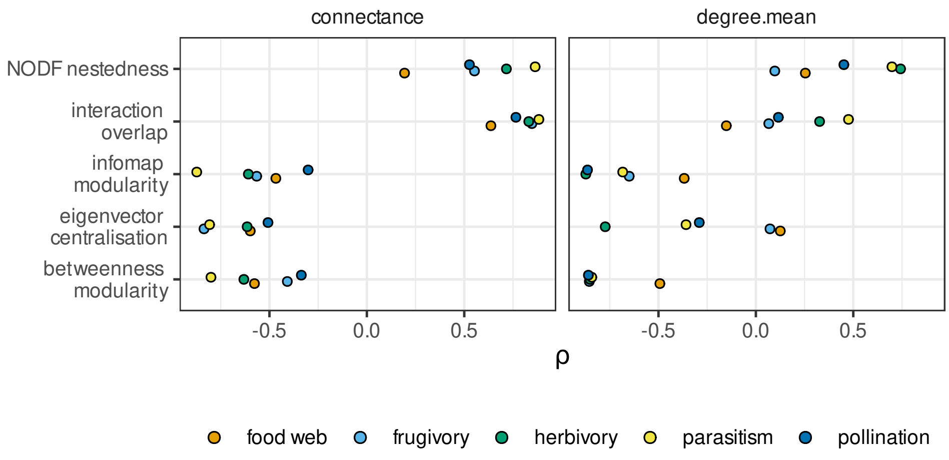


Fig. S3: Spearman correlations between network metrics obtained and connectance (left panel) and degree distribution average (right panel), for networks of each interaction type.

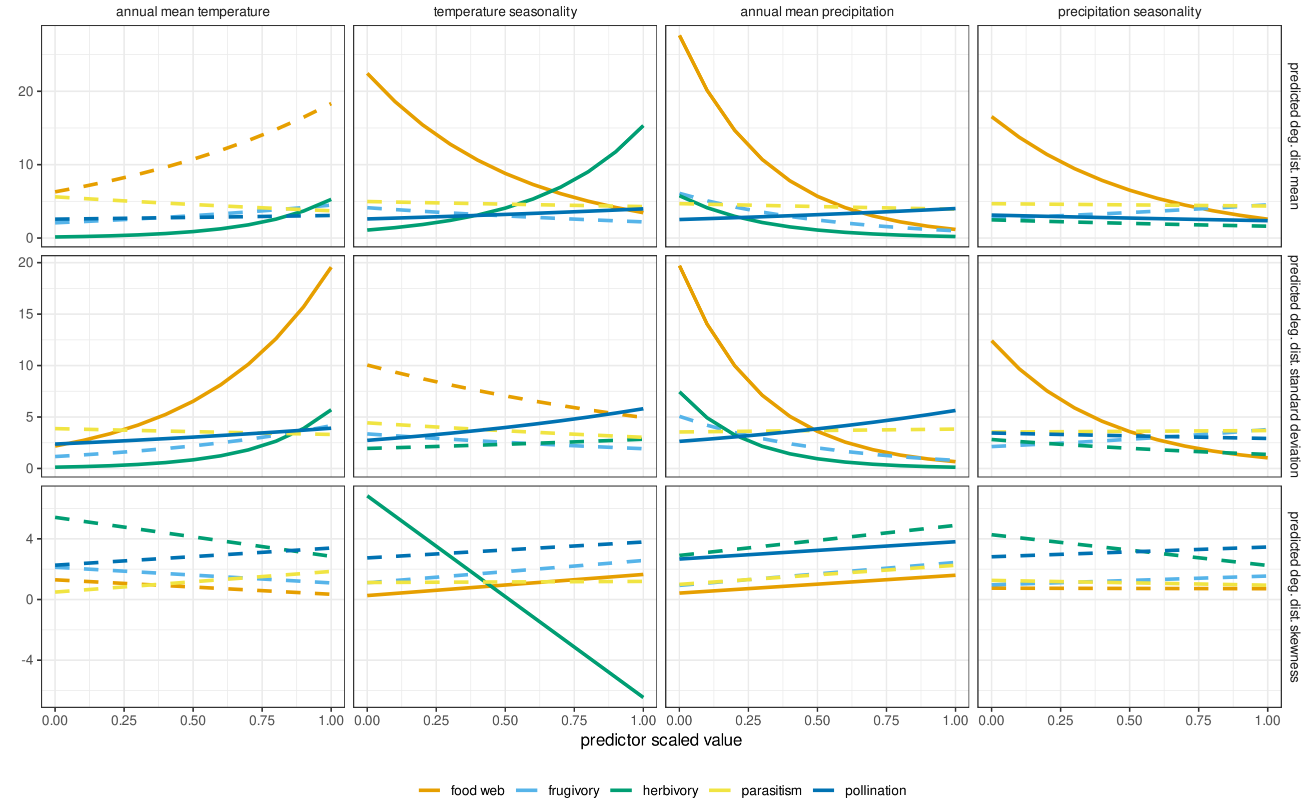


Fig. S4: As Fig. 5 of main text, for models without study as random factor.

### Regression tables

In the following tables, note that the mean and standard deviation regressions had the response variable log-transformed; coefficients are in the original log scale. Skewness was not log-transformed.

#### Response: degree distribution average

##### Food webs

| effect | component | group | term | estimate | std.error | statistic | p.value |
|---|---|---|---|---|---|---|---|
| fixed | cond | | (Intercept) | 3.41 | 0.58 | 5.85 | 0.00 |
| fixed | cond | | annual_mean_temp | -0.26 | 0.82 | -0.32 | 0.75 |
| fixed | cond | | temp_seasonality | -1.44 | 0.65 | -2.20 | 0.03 |
| fixed | cond | | annual_mean_prec | -1.58 | 0.34 | -4.58 | 0.00 |
| fixed | cond | | prec_seasonality | 0.03 | 0.36 | 0.07 | 0.94 |
| ran_pars | cond | study_common | sd__(Intercept) | 0.58 | | | |
| ran_pars | cond | Residual | sd__Observation | 0.32 | | | |

##### Frugivory

| effect | component | group | term | estimate | std.error | statistic | p.value |
|---|---|---|---|---|---|---|---|
| fixed | cond | | (Intercept) | 0.98 | 1.05 | 0.93 | 0.35 |
| fixed | cond | | annual_mean_temp | 0.77 | 1.35 | 0.57 | 0.57 |
| fixed | cond | | temp_seasonality | -0.63 | 2.10 | -0.30 | 0.76 |
| fixed | cond | | annual_mean_prec | -1.82 | 1.36 | -1.34 | 0.18 |
| fixed | cond | | prec_seasonality | 0.53 | 0.54 | 0.98 | 0.33 |
| ran_pars | cond | study_common | sd__(Intercept) | 0.00 | | | |
| ran_pars | cond | Residual | sd__Observation | 0.14 | | | |

##### Herbivory

| effect | component | group | term | estimate | std.error | statistic | p.value |
|---|---|---|---|---|---|---|---|
| fixed | cond | | (Intercept) | 0.40 | 0.77 | 0.52 | 0.61 |
| fixed | cond | | annual_mean_temp | -0.24 | 1.06 | -0.22 | 0.82 |
| fixed | cond | | temp_seasonality | 1.07 | 1.22 | 0.87 | 0.38 |
| fixed | cond | | annual_mean_prec | -0.58 | 0.70 | -0.83 | 0.40 |
| fixed | cond | | prec_seasonality | 1.23 | 0.69 | 1.79 | 0.07 |
| ran_pars | cond | study_common | sd__(Intercept) | 0.26 | | | |
| ran_pars | cond | Residual | sd__Observation | 0.09 | | | |

##### Parasitism

| effect | component | group | term | estimate | std.error | statistic | p.value |
|---|---|---|---|---|---|---|---|
| fixed | cond | | (Intercept) | 1.85 | 0.77 | 2.40 | 0.02 |
| fixed | cond | | annual_mean_temp | -0.41 | 0.55 | -0.75 | 0.46 |
| fixed | cond | | temp_seasonality | -0.15 | 0.75 | -0.20 | 0.84 |
| fixed | cond | | annual_mean_prec | -0.18 | 1.20 | -0.15 | 0.88 |
| fixed | cond | | prec_seasonality | -0.07 | 0.32 | -0.22 | 0.83 |
| ran_pars | cond | study_common | sd__(Intercept) | 0.00 | | | |
| ran_pars | cond | Residual | sd__Observation | 0.35 | | | |

##### Pollination

| effect | component | group | term | estimate | std.error | statistic | p.value |
|---|---|---|---|---|---|---|---|
| fixed | cond | | (Intercept) | 1.04 | 0.28 | 3.75 | 0.00 |
| fixed | cond | | annual_mean_temp | 0.09 | 0.32 | 0.27 | 0.78 |
| fixed | cond | | temp_seasonality | 0.14 | 0.24 | 0.60 | 0.55 |
| fixed | cond | | annual_mean_prec | 0.09 | 0.14 | 0.66 | 0.51 |
| fixed | cond | | prec_seasonality | -0.11 | 0.17 | -0.65 | 0.52 |
| ran_pars | cond | study_common | sd__(Intercept) | 0.34 | | | |
| ran_pars | cond | Residual | sd__Observation | 0.16 | | | |

#### Response: degree distribution standard deviation

##### Food webs

| effect | component | group | term | estimate | std.error | statistic | p.value |
|---|---|---|---|---|---|---|---|
| fixed | cond | | (Intercept) | 3.30 | 0.53 | 6.17 | 0.00 |
| fixed | cond | | annual_mean_temp | -0.55 | 0.75 | -0.72 | 0.47 |
| fixed | cond | | temp_seasonality | -1.98 | 0.60 | -3.32 | 0.00 |
| fixed | cond | | annual_mean_prec | -1.52 | 0.31 | -4.86 | 0.00 |
| fixed | cond | | prec_seasonality | -0.07 | 0.33 | -0.20 | 0.84 |
| ran_pars | cond | study_common | sd__(Intercept) | 0.53 | | | |
| ran_pars | cond | Residual | sd__Observation | 0.29 | | | |

##### Frugivory

| effect | component | group | term | estimate | std.error | statistic | p.value |
|---|---|---|---|---|---|---|---|
| fixed | cond | | (Intercept) | 0.37 | 1.01 | 0.36 | 0.72 |
| fixed | cond | | annual_mean_temp | 1.28 | 1.29 | 0.99 | 0.32 |
| fixed | cond | | temp_seasonality | -0.56 | 2.00 | -0.28 | 0.78 |
| fixed | cond | | annual_mean_prec | -1.86 | 1.30 | -1.44 | 0.15 |
| fixed | cond | | prec_seasonality | 0.58 | 0.52 | 1.12 | 0.26 |
| ran_pars | cond | study_common | sd__(Intercept) | 0.00 | | | |
| ran_pars | cond | Residual | sd__Observation | 0.13 | | | |

##### Herbivory

| effect | component | group | term | estimate | std.error | statistic | p.value |
|---|---|---|---|---|---|---|---|
| fixed | cond | | (Intercept) | -0.67 | 1.04 | -0.64 | 0.52 |
| fixed | cond | | annual_mean_temp | 3.83 | 1.92 | 2.00 | 0.05 |
| fixed | cond | | temp_seasonality | 0.38 | 0.58 | 0.66 | 0.51 |
| fixed | cond | | annual_mean_prec | -4.14 | 0.81 | -5.10 | 0.00 |
| fixed | cond | | prec_seasonality | -0.72 | 0.72 | -1.00 | 0.31 |
| ran_pars | cond | study_common | sd__(Intercept) | 0.00 | | | |
| ran_pars | cond | Residual | sd__Observation | 0.24 | | | |

##### Parasitism

| effect | component | group | term | estimate | std.error | statistic | p.value |
|---|---|---|---|---|---|---|---|
| fixed | cond | | (Intercept) | 1.54 | 0.73 | 2.11 | 0.03 |
| fixed | cond | | annual_mean_temp | -0.16 | 0.52 | -0.30 | 0.76 |
| fixed | cond | | temp_seasonality | -0.38 | 0.71 | -0.54 | 0.59 |
| fixed | cond | | annual_mean_prec | 0.08 | 1.13 | 0.07 | 0.95 |
| fixed | cond | | prec_seasonality | 0.04 | 0.31 | 0.13 | 0.89 |
| ran_pars | cond | study_common | sd__(Intercept) | 0.00 | | | |
| ran_pars | cond | Residual | sd__Observation | 0.33 | | | |

##### Pollination

| effect | component | group | term | estimate | std.error | statistic | p.value |
|---|---|---|---|---|---|---|---|
| fixed | cond | | (Intercept) | 0.66 | 0.39 | 1.71 | 0.09 |
| fixed | cond | | annual_mean_temp | 0.80 | 0.45 | 1.78 | 0.07 |
| fixed | cond | | temp_seasonality | 0.69 | 0.36 | 1.93 | 0.05 |
| fixed | cond | | annual_mean_prec | 0.07 | 0.21 | 0.33 | 0.74 |
| fixed | cond | | prec_seasonality | -0.30 | 0.24 | -1.24 | 0.22 |
| ran_pars | cond | study_common | sd__(Intercept) | 0.41 | | | |
| ran_pars | cond | Residual | sd__Observation | 0.26 | | | |

#### Response: degree distribution skewness

##### Food webs

| effect | component | group | term | estimate | std.error | statistic | p.value |
|---|---|---|---|---|---|---|---|
| fixed | cond | | (Intercept) | 3.47 | 0.84 | 4.12 | 0.00 |
| fixed | cond | | annual_mean_temp | -3.65 | 1.17 | -3.13 | 0.00 |
| fixed | cond | | temp_seasonality | -2.68 | 0.95 | -2.82 | 0.00 |
| fixed | cond | | annual_mean_prec | 1.19 | 0.49 | 2.43 | 0.02 |
| fixed | cond | | prec_seasonality | 0.68 | 0.51 | 1.32 | 0.19 |
| ran_pars | cond | study_common | sd__(Intercept) | 0.83 | | | |
| ran_pars | cond | Residual | sd__Observation | 0.45 | | | |

##### Frugivory

| effect | component | group | term | estimate | std.error | statistic | p.value |
|---|---|---|---|---|---|---|---|
| fixed | cond | | (Intercept) | 1.12 | 3.54 | 0.32 | 0.75 |
| fixed | cond | | annual_mean_temp | -1.03 | 4.53 | -0.23 | 0.82 |
| fixed | cond | | temp_seasonality | 1.47 | 7.05 | 0.21 | 0.83 |
| fixed | cond | | annual_mean_prec | 1.51 | 4.56 | 0.33 | 0.74 |
| fixed | cond | | prec_seasonality | 0.58 | 1.82 | 0.32 | 0.75 |
| ran_pars | cond | study_common | sd__(Intercept) | 0.00 | | | |
| ran_pars | cond | Residual | sd__Observation | 0.47 | | | |

##### Herbivory

| effect | component | group | term | estimate | std.error | statistic | p.value |
|---|---|---|---|---|---|---|---|
| fixed | cond | | (Intercept) | 8.92 | 4.39 | 2.03 | 0.04 |
| fixed | cond | | annual_mean_temp | -2.57 | 8.10 | -0.32 | 0.75 |
| fixed | cond | | temp_seasonality | -13.30 | 2.44 | -5.46 | 0.00 |
| fixed | cond | | annual_mean_prec | 1.98 | 3.43 | 0.58 | 0.56 |
| fixed | cond | | prec_seasonality | -2.03 | 3.04 | -0.67 | 0.51 |
| ran_pars | cond | study_common | sd__(Intercept) | 0.00 | | | |
| ran_pars | cond | Residual | sd__Observation | 1.00 | | | |

##### Parasitism

| effect | component | group | term | estimate | std.error | statistic | p.value |
|---|---|---|---|---|---|---|---|
| fixed | cond | | (Intercept) | 0.38 | 1.50 | 0.26 | 0.80 |
| fixed | cond | | annual_mean_temp | 1.36 | 1.08 | 1.26 | 0.21 |
| fixed | cond | | temp_seasonality | 0.07 | 1.45 | 0.05 | 0.96 |
| fixed | cond | | annual_mean_prec | 1.26 | 2.33 | 0.54 | 0.59 |
| fixed | cond | | prec_seasonality | -0.32 | 0.63 | -0.50 | 0.61 |
| ran_pars | cond | study_common | sd__(Intercept) | 0.00 | | | |
| ran_pars | cond | Residual | sd__Observation | 0.68 | | | |

##### Pollination

| effect | component | group | term | estimate | std.error | statistic | p.value |
|---|---|---|---|---|---|---|---|
| fixed | cond | | (Intercept) | 2.18 | 1.08 | 2.03 | 0.04 |
| fixed | cond | | annual_mean_temp | 1.29 | 1.23 | 1.05 | 0.29 |
| fixed | cond | | temp_seasonality | 1.32 | 1.12 | 1.18 | 0.24 |
| fixed | cond | | annual_mean_prec | -0.39 | 0.66 | -0.58 | 0.56 |
| fixed | cond | | prec_seasonality | -0.27 | 0.73 | -0.37 | 0.71 |
| ran_pars | cond | study_common | sd__(Intercept) | 0.92 | | | |
| ran_pars | cond | Residual | sd__Observation | 0.93 | | | |
